# Exploring the microbial diversity in oil-contaminated mangrove sediments using 16S rRNA metagenomics

**DOI:** 10.1101/735290

**Authors:** Mahdi Ghanbari, Mansooreh Jami, Mohsen Shahriari Moghadam, Konrad J. Domig

## Abstract

With an area of 390 hectares, the mangrove forests of Nayband Bay are the widest mangrove communities above 27 degrees latitude in the Persian Gulf. They are the last dense, extensive structures of these ecosystems found in the north-west Indian Ocean. Growing industrial activities near the Nayband Bay and the consequent marine pollution has resulted in damage and threatened different marine habitats in recent years. To date, a comprehensive description of the microbial life in the mangrove ecosystem that has been exposed to oil contamination is lacking. This information could significantly contribute to a better overview of the function and resilience of the ecosystem. This work represents a first effort to better understand the Nayband Bay mangrove microbiology by applying 16S rRNA metagenomics. A total of 65,408 readings from the V3–V4 16S rRNA gene regions were obtained from 24 sediment samples, each measuring 440 bp. Most sequences belonged to members of the Proteobacteria phylum (mainly γ-Proteobacteria); however, members of the Bacteroidetes phylum (mainly Flavobacteriia) were also well represented in the samples. We discovered that the community of this ecosystem strongly exhibit typical structures of oil-contaminated marine environments. This is likely due to the growing industrial activity in the area and its consequent marine polluting effects. The use of practicable and applicable bioremediation protocols for habitat restoration in this valuable area is needed.

## Introduction

Mangrove ecosystems constitute a large portion (60–70%) of the coastline in the tropical and subtropical regions of Earth [1]. Iranian mangrove forests occur between longitude 25’ 190 and 27’ 84, in the northern part of the Persian Gulf and Oman Sea [2]. With an area of 390 hectares, the mangrove forests of Nayband Bay are the widest mangrove communities above 27 degrees latitude in the Persian Gulf. They are the last dense, extensive structures of these ecosystems in the north-west Indian Ocean [2]. These mangroves support more than 84 species of waterbirds and large populations of migratory birds and is one of the most important feeding grounds and nesting sites for hawksbill and green sea turtles [2]. Despite its ecological importance in the Persian Gulf, Nayband Bay is surrounded by oil and gas facilities of the Pars Special Economic Energy Zone (PSEEZ) in the north-west. Growing industrial activity in the area and its consequent marine polluting effects has resulted in damage and has threatened marine habitats in the Nayband Bay in recent years [3, 2]. Several studies have shown that contamination with petroleum hydrocarbons affects the structural and functional diversity of bacterial populations in the contaminated soils and sediments [4-6], which may have large ecological implications. Although there are some reports on the microbiology of the mangrove environment in Nayband Bay [3, 7-9], almost all of the studies have concentrated on the culture-dependent analysis of polycyclic aromatic hydrocarbon (PAH)-degrading bacteria from this environment. To date, a comprehensive description of the microbial life in this mangrove ecosystem that has been exposed to oil pollution is lacking.

In this study, we used a 16S rRNA amplicon sequencing approach to investigate the composition of the bacterial communities in Nayband Bay mangrove sediments. With the recent advances in high-throughput DNA sequencing techniques, 16S rRNA gene amplicon sequencing has frequently been used to study the microbial ecology of a wide range of environments [6, 10, 11]. To the best of our knowledge, the present study is the first report that uses 16S rRNA-metagenomics to study the microbial communities present in Nayband Bay mangrove sediments. Importantly, the findings of this study could significantly contribute to a better overview of the functioning and resilience of the ecosystem.

## Materials and methods

### Sample collection and DNA extraction

For this study, twenty-eight sediment samples were obtained randomly using a sediment core (7 cm diameter and 30 cm depth) from seven sites (each site 100 m^2^) in the mangrove forests of Nayband Bay (27° 27.519’ N52° 40.545’ E) in the Persian Gulf during low tide (Fig. 1). Physicochemical properties of environmental factors of the sampling area (pH□=□8; temperature□=□30°C; salinity□=□40 ppt) were measured using an HQ40d Multimeter (HACH Colorado, USA). Equal amounts of each sediment sample were mixed together thoroughly to form a composite sample. Three 0.3 g aliquots of homogenised sediment were subjected to DNA extraction using the Power Soil DNA Isolation kit (MoBio® Laboratories Inc., Carlsbad, CA, USA). DNA was visualised on a 0.8% agarose gel and quantified using a Qubit 2.0 fluorometer (Invitrogen, Carlsbad, CA, USA). DNA from all samples were pooled together and the DNA was concentrated to a final volume of 50 µl. Extracted DNA was stored at −80 °C.

**Fig. 1.**
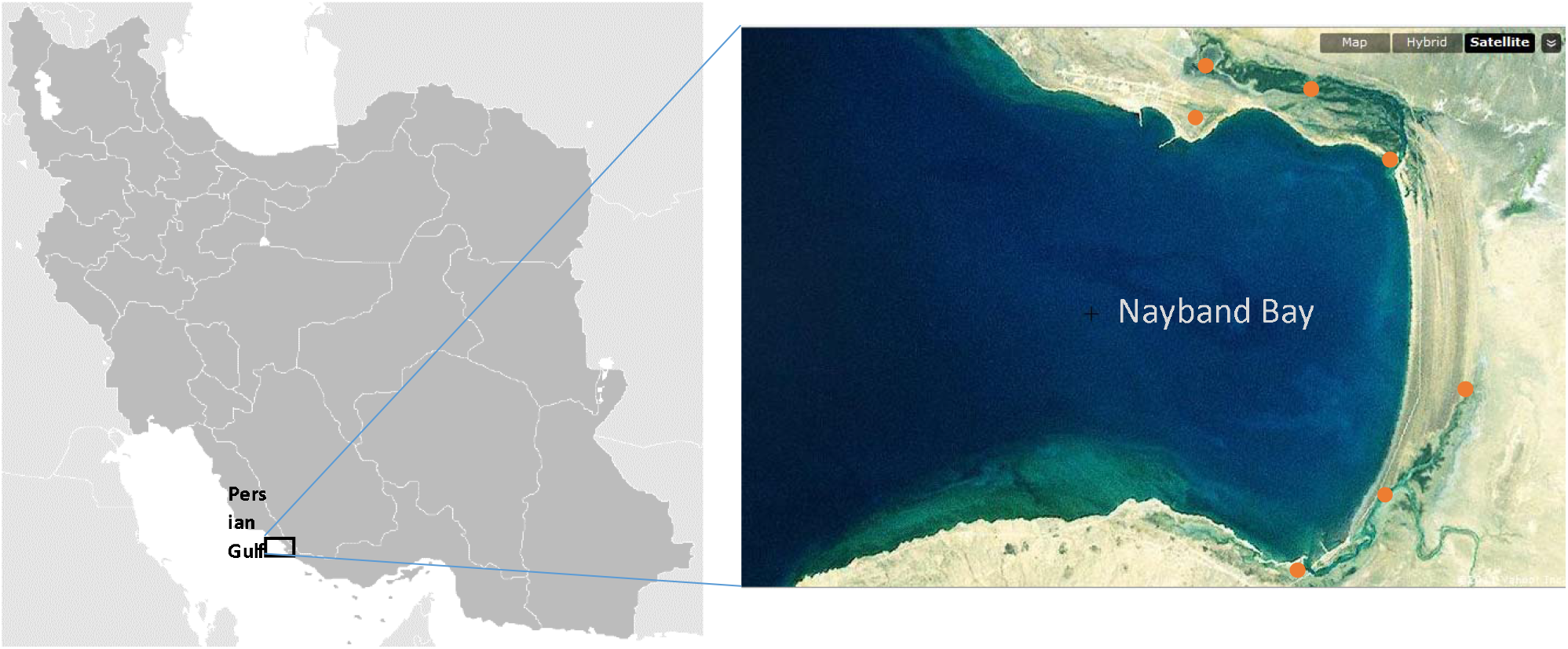
Map of the study area and location of the mangrove sampling sites at Nayband Bay.

### Molecular analysis and bioinformatics

Detailed information on the 16Sbased target enrichment, PCR conditions, 2 × 300 bp sequencing process (Illumina, MiSeq V3), and data analysis can be found in our previous articles [12, 13]. Briefly, a two-step PCR amplification was performed on the samples, negative extraction controls, and reagent blanks in duplicate using Illumina forward overhang adapter sequence [5□-TCGTCGGCAGCGTCAGA TGTGTATAAGAGACAG-(locus-specific sequence)-3’] and reverse overhang adapter sequence [5□-GTCTCGTGGGCTCGGAGATGTGTATAAGAGACAG-(locus-specific sequence)-3’] along with the locus-specific sequence using S-D-Bact-0341-b-S-17 (5’-CCTACGGGNGGCWGCAG-3’) as forward primer and S-D-Bact-0785-a-A21 (5’-GACTACHVGGGTATCTAATCC-3’) as reverse primer, followed by adding the Illumina sequencing adapters. The quality of the amplified library was evaluated fluorometrically (Qubit 3, Invitrogen), and purity was assessed via the 260/280 and 260/230 absorbance ratios, as determined via spectrophotometry (NanoDrop® ND-1000). The samples were sent for DNA sequencing to the MicroSynth AG sequencing facility (Balgach, Switzerland). Paired-end sequencing libraries (2 × 300 bp) were prepared using the Illumina Nextera XT Library Preparation Kit (Illumina Inc., San Diego, CA) and sequenced on the Illumina MiSeq platform using V3 chemistry according to the manufacturer’s instructions. All analyses were performed with custom Bash and R, and Perl scripts, building off of established concepts and utilizing existing algorithms and toolkits such UPARSE [14] and QIIME v 1.9.1 [15] pipelines as mentioned in our previous reports [16, 12]. The raw sequence data are available in the Sequence Read Archive at the National Center for Biotechnology Information (BioProject accession no. PRJNA331081; SRA accession no. SRP079708).

## Results and discussion

To date, extensive research has focused on oil bioremediation using pure cultures or mixed bacterial consortia isolated from oil spilled mangrove soils. However, few studies have been conducted on the different bacterial communities and diversity in mangrove sediments exposed to crude oil contamination. The aim of this study was to assess the bacterial community structure and composition in sediments of the Nayband Bay mangrove forests that had been negatively affected by oil pollution as a result of development of the South Pars oil field in the Assaluyeh region [3, 2]. To our knowledge, this is the first description of the bacterial community found in Iranian mangrove sites using an NGS approach. A total of 65,408 readings from the V3–V4 16S rRNA gene regions were obtained from a mix of 24 sediment samples, each measuring 440 bp and were assigned to bacterial OTU’s at a minimum sequence homology of 97%. Fig. 2 and Fig. S1 illustrate the abundance of major (>1%) phyla, classes, families and genera observed in the mangrove soil samples. Details about the microbial taxon composition, diversity, frequencies and prevalence are given in supplementary data 1. Bacteria representing three phyla were found in the samples. Most sequences belonged to members of the Proteobacteria phylum (mean sequence abundance 69%); however, members of the Bacteroidetes phylum (30%) was also well represented in the samples. Firmicutes, with an abundance of <1%, was the third most prevalent phylum found in the samples (Fig. S1). The most abundant classes recorded, in terms of mean sequence abundance, were γ-Proteobacteria (69%) and Flavobacteriia (30%), with the rest of the classes present in lower sequence abundance (<1%). Although no similar studies have been conducted on the Nayband Bay mangrove forests, surveys on other tropical mangrove sediment bacterial communities suggest that Proteobacteria, Bacteroidetes and Firmicutes are numerically the most dominant phyla, even after exposure to oil contamination [6, 1]. Consistent with previous studies, the present study also found Proteobacteria and Bacteroidetes to be the most abundant in mangroves, although Firmicutes abundance was lower in comparison with the previous reports. In fact, the dominance of sulfate-reducing bacteria (which are mainly from Proteobacteria) in the mangrove environment is not surprising, since mangrove soils are anaerobic environments rich in sulfate and organic matter [6].

**Fig. 2.**
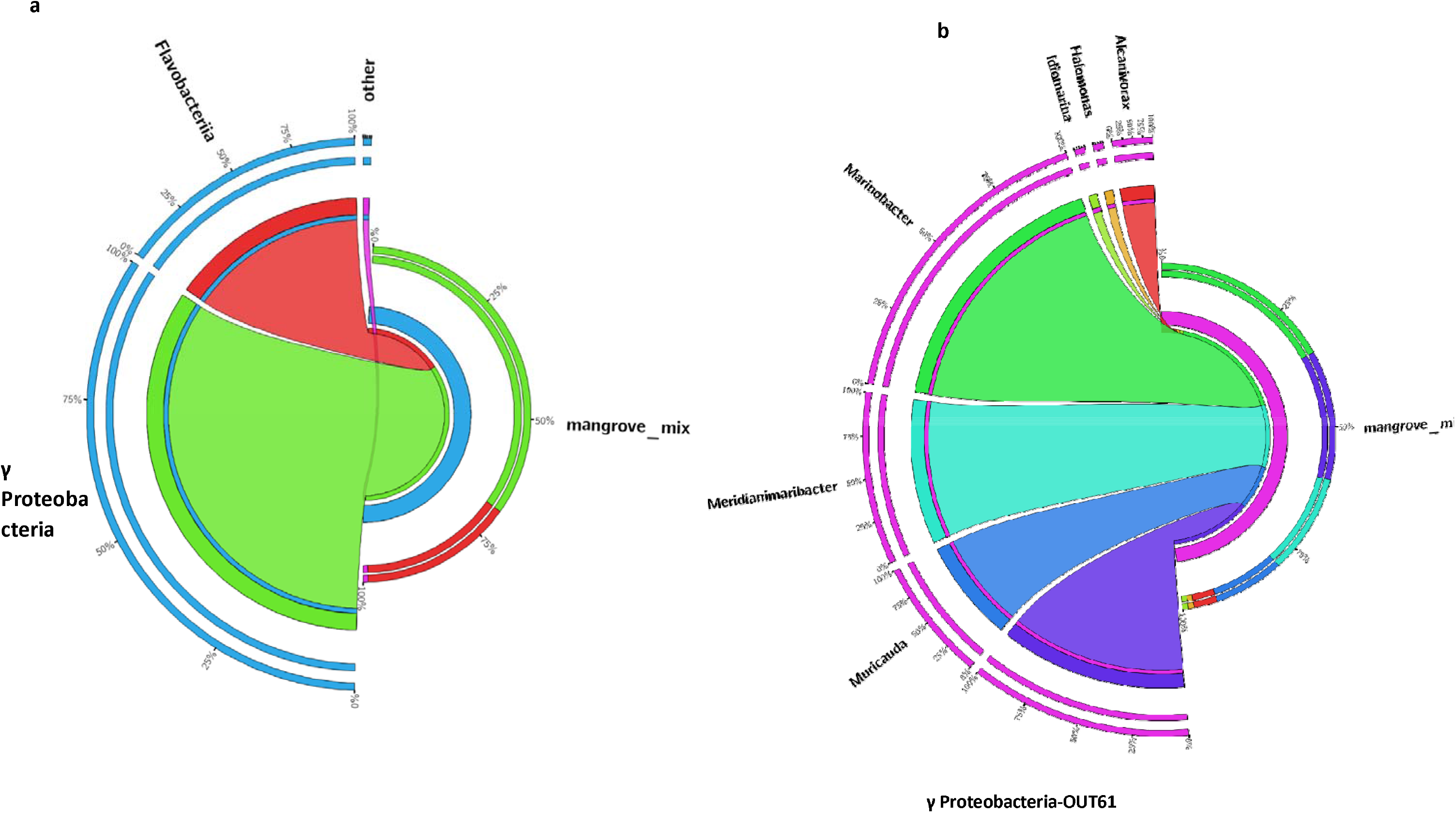
Circular representation of the microbial communities in mangrove sediment samples at the class (a) and genus (b) level. Taxa with a relative abundance lower than 1% are not shown. The circular representation for other taxonomic levels can be found in Fig. S1.

The proteobacterial community of the Nayband mangrove sediments was dominated by members of the γ-Proteobacteria class, mainly members of the Alteromonadales and Oceanospirillales orders (the so-called *Marinobacter hydrocarbonoclasticus*) (supplementary data 1). Interestingly, the high presence of these groups of bacteria in marine and mangrove sediments has been significantly linked to crude oil contamination [6, 17-19] and are of major importance for the biodegradation of marine crude oil contaminants [20, 21, 18]. For instance, dos Santos et al. [6] reported a notable increase in the abundance of Alteromonadales and Oceanospirillales in oil-treated samples (with 2-5% oil). In our study, *Marinobacter* (31%), followed by *Idiomarina* (9%), was the most abundant of the Alteromonadales order within the mangrove sediment samples (Fig. 2, supplementary data 1). These genera have already been described in several studies as being present in hypersaline waters and areas contaminated with oil, including mangroves [22, 6, 23]. The high abundance of these genera in the present study (*Marinobacter*, followed by *Idiomarina)* may highlight their role as potential proxies for the biomonitoring of petroleum hydrocarbons in this ecosystem. The *Haliea* genus from the Alteromonadales order only represented a small percentage of the community (0.3%). The members of this genus have been isolated before from the marine environment [24, 6]. The low abundance of this genus in comparison to other genera of the Alteromonadales order suggests that these organisms are sensitive to oil. Similarly, a recent study research detected a significant decline in the abundance of *Haliea* genes after exposure to 2-5% oil for over 23 days [6].

In the Oceanospirillales order, the genus that displayed the greatest abundance in the present study was *Alcanivorax* (4%), followed by *Halomonas*, which was poorly represented at 1% (Fig. 2, supplementary data 1). Both genera have been reported previously in relation to the biodegradation of crude oil contamination in the marine environment [25, 19]. *Alcanivorax* species are alkane-degrading marine bacteria that are able to propagate and become dominant in crude oil-containing seawater supplemented with nitrogen and phosphorus [17, 19, 23]; this characteristic has been observed in the deep water oil plume at the Gulf spill site as well as in laboratory enrichments and in the field after exposure to hydrocarbons [24].

Bacteroidetes was the second most abundant phylum in the present dataset (Fig. S1, supplementary data 1). Almost all the Bacteroidetes sequences were assigned to the Flavobacteriia class. This class of bacteria is widespread in various aquatic environments, including marine areas and mangrove sediments, and probably plays a major role in the turnover of organic matter, since they are able to degrade natural polymers (e.g., starch, cellulose, or gelatine) [26-28]. The Flavobacteriaceae family is one of the most widely expanding families in the *Bacteroidetes* phylum; more than 50 genera have been described, most of them in recent years [29]. Most of the recently described genera of the Flavobacteriaceae, e.g., *Meridianimaribacter, Muricauda, Psychroserpens, Gelidibacter*, and *Cellulophaga*, were isolated from marine environments, demonstrating that this is an important habitat for members of the Flavobacteriia class [30, 31, 29, 28, 26]. While there is a little information, some members of the Flavobacteriaceae family have exhibited detectable PAHs degradability and may serve as efficient degraders of crude oil contamination of mangrove ecosystems [27, 32, 30, 11, 26]. Both members of this family that were present in the current study, *Meridianimaribacter* (17%) and *Muricauda* (12%), have been detected previously in marine samples and mangrove sediments [29, 31, 28, 30].

## Conclusion

The evaluation and monitoring of biological activity in valuable ecosystems such as the mangrove forests includes understanding the effects of the biotic and abiotic factors, such as the effects of the oil spills on the composition and function of the indigenous microbiome. Using 16S rRNA metagenomics, we conducted an analysis of the microbiota membership and structure to obtain a comprehensive understanding of the microbial life in a mangrove area located in Nayband Bay, Iran, where the South Pars Gas Complex (SPGC) and the Assaluyeh harbour have been established and enlarged. Growing industrial activity in the area and its consequent marine pollution make this marine environment highly susceptible to an ecological disaster caused by an oil spill. The results show that, in addition to the presence of many sulfate-reducing bacteria, the community of this ecosystem displays characteristics typical of the structure of oil-contaminated marine environments. This finding is likely due to the growing industrial activity in the area and its consequent marine polluting effects (e.g., SPGC). Therefore, the use of practicable and applicable bioremediation and conservation protocols for habitat restoration in this valuable area is necessary. Although, due to technical limitations, the low number of samples had to be pooled and investigated, the methodology applied in this survey provides a first look at the microbial community composition that underlies the biogeochemical transformations occurring in this environment. As the structure and composition of microbial communities in mangrove forests are influenced by different biotic and abiotic factors, future work in this area should focus on a complete description of the potential metabolic traits of these organisms and their dynamics throughout the year. Important advances will be achieved by applying whole metagenome shotgun sequencing in mangrove sediments collected from different sites.

## Acknowledgments

We thank Dr. Behrouz Abtahi, Hadi Shahraki (UOZ) and Dr. Manuel Kraler (BOKU), for their outstanding (technical) assistance to organize collecting and processing of the samples.

