## Supplementary material for "Exploring the microbial diversity in oil-contaminated mangrove sediments using 16S rRNA metagenomics": Figure S1

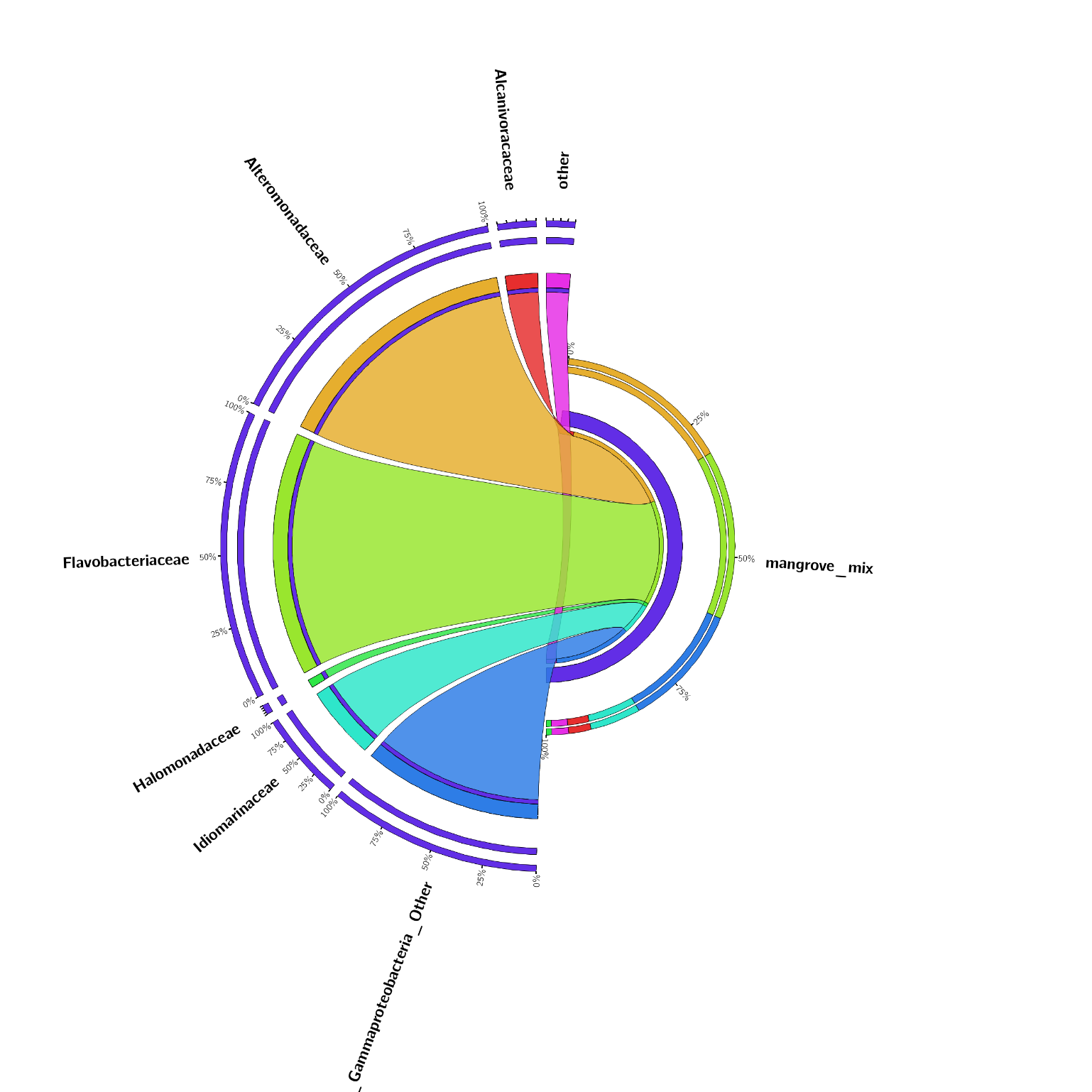

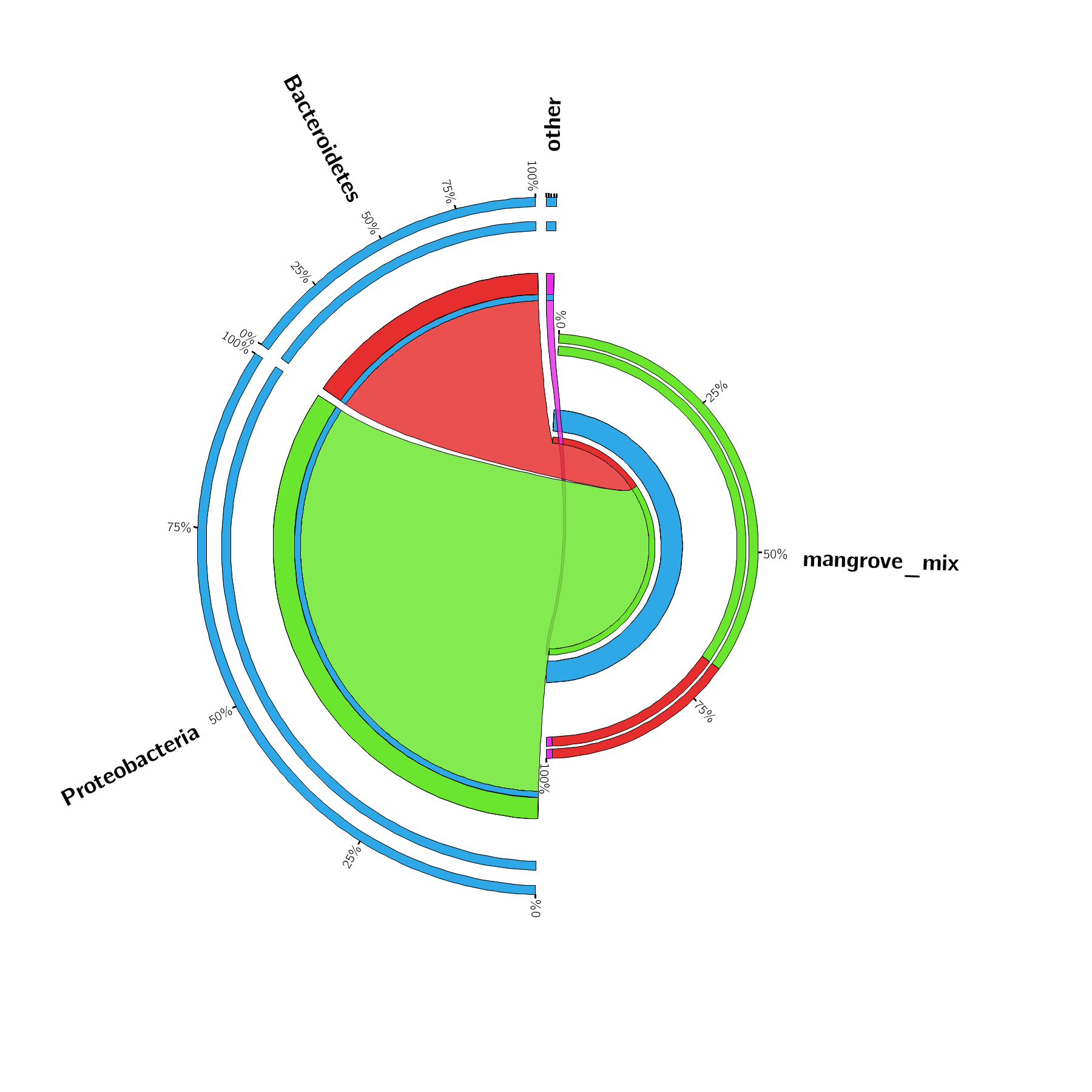

Fig S1. Circular representation of microbial communities in mangrove sediment samples at phylum (a) and family (b). Taxa with relative abundance lower than 1% were not shown.
